# Highly efficient generation of blastocyst-like structures from mouse totipotent blastomere-like cells

**DOI:** 10.1101/2022.03.01.481880

**Authors:** Pengfei Zhang, Xuzhao Zhai, Boyan Huang, Shu Sun, WenJing Wang, Man Zhang

## Abstract

Mammalian embryogenesis begins with a totipotent zygote. Early embryogenesis can be recapitulated by aggregated extended pluripotent stem cells (EPSCs) in a 3D culture system. However, the efficiency of generating blastoids is low and whether other reported totipotent stem cells retained a similar capacity remains unknown. Here we show that spliceosomal repressed totipotent blastomere-like cells (TBLCs) form blastocyst-like structures when aggregated in 3D microwells with around 80% efficiency. TBLC-blastoids resemble blastocyst in morphology and cell-lineage allocation, show similar transcriptional profile with natural blastocyst and contain more TE cells and fewer undefined intermediate cells compared to blastoids from EPSCs. Moreover, TBLC-blastoids can develop beyond the implantation stage *in vitro* and induce decidualization after transferred into uterus. In summary, we supply an alternative cell type to generate ameliorated blastoids highly efficiently for studying early mouse embryogenesis.

## INTRODUCTION

Mammalian embryogenesis begins with a totipotent zygote which is formed by the fusion of gametes (McGrath and Solter, 1984). After fertilization, the single totipotent zygote divides and initiates the first cell lineage segregation at 8-cell stage, eventually forms a blastocyst before implantation which contains the earliest three cell lineages, inner cell mass (ICM)/epiblast (Epi), trophectoderm (TE) and primitive endoderm (PrE) (Rossant and Tam, 2009; Saiz and Plusa, 2013). Cells of epiblast will develop into the embryo proper, whereas cells of the TE and PrE will give rise to the placenta and yolk sac (Rossant, 2008; Rossant and Tam, 2009). Counterpart cell lines, namely embryonic stem cell (ESC) (Bradley et al., 1984; Evans and Kaufman, 1981), trophoblast stem cell (TSC) (Tanaka et al., 1998) and extraembryonic endoderm stem cell (XEN) (Kunath et al., 2005; Niakan et al., 2013) or primitive endoderm stem cell (PrESCs) (Ohinata et al., 2022) can be derived from ICM/epiblast, TE and PrE respectively when cultured blastocysts *in vitro*. These three types of cell lines are characteristic of the early cell lineages and cells from them can incorporate into the embryo development in compliance with their defined developmental potential after being injected into the blastocysts (Bradley et al., 1984; Kruithof-de Julio et al., 2011; Kunath et al., 2005; Ohinata et al., 2022; Tanaka et al., 1998). Interestingly, previous work showed that aggregation of ESCs and TSCs without or with XEN cells self-assemble to generate embryo-like structures whose morphogenesis, constituent cell-types and transcriptional profile resemble blastocysts or early post-implantation natural embryos, which were referred as ETS-blastoids or ETS-embryos respectively (Harrison et al., 2017; Rivron et al., 2018; Sozen et al., 2018). Moreover, Li et.al, reported that extended pluripotent stem (EPS) cells which can contribute to both embryonic and extraembryonic lineages *in vitro* and *in vivo* (Yang et al., 2017a; Yang et al., 2017b) are capable of generation of blastoids (EPS-blastoids) in the medium contains FGF4, BMP4, CHIR99021 and A83-01 on their own (Li et al., 2019; Sozen et al., 2019). These EPS-blastoids recapitulate pre-implantation development process and that of early post-implantation after cultured *in vitro* culture (IVC) medium (Bedzhov et al., 2014), including compaction, polarization, rosette formation and lumenogenesis. Moreover, EPS-blastoids are capable of implantation, triggering decidualization *in vivo*. Thus, this single cell type derived blastoids system could be used as a tractable, highly malleable and consecutive *in vitro* model for studying preimplantation and early postimplantation embryogenesis. However, the average efficiency of EPS-blastoids generation is only 15% and a considerable amount (60%) of uncommitted intermediated cells which are closely correlated to mesoderm exist within the EPS-blastoids (Li et al., 2019; Posfai et al., 2021). These impede the usage of EPS-blastoids to study early embryogenesis. Although EPS cells express some molecular features of totipotent blastomeres, their transcriptome is close to E4.5 or E5.5 Epi (Posfai et al., 2021; Shen et al., 2021; Yang et al., 2017a). Recent work showed EPS cells have barely extraembryonic development potential in chimeras and in TSC culture condition in vitro (Posfai et al., 2021). Recently, Shen et.al reported that spliceosomal repression can reprogram pluripotent mouse ESCs to a totipotent state. Moreover, stable *in vitro* culture of totipotent cells is achieved by treated ESCs with splicing inhibitor pladienolide B (PlaB). These totipotent cells which they called totipotent blastomere-like cells (TBLCs) are comparable at molecular levels with 2- and 4-cell blastomeres and show robust bidirectional developmental capability to generate both embryonic and extraembryonic cell lineages in chimeras (Shen et al., 2021). However, whether totipotent TBLCs retain the ability to form blastoids *in vitro* is unknown. Here we show that TBLCs form blastoids in the 3D differentiation culture system highly efficiently.

## RESULTS

### TBLCs form blastoids *in vitro* in a 3D differentaition system

Recently, Du’s laboratory reported that spliceosomal repression reprograms pluripotent mouse ESCs to a totipotent state, in which their transcriptome is close to 2- and 4-cell blastomeres, which they called them totipotent blastomere-like cells (TBLCs). To obtain TBLCs, we cultured mCMG (Huang et al., 2022) ESCs in ES medium plus 2.5 nM PlaB on feeders for over 6 passages, and named them as mCMG-TBLCs (Figure S1A). Consistent with previous report (Shen et al., 2021), transcriptome analysis showed that passage 12 mCMG-TBLCs upregulated 2C or 4C-speicific genes, *Zscan4c/4d, Zfp352, Mmp19, Snai1, Btg1*, etc. and downregulated pluripotent mRNAs, *Zfp42, Tet1*, *Sox2, Pou5f1*(*Oct4*), etc (Figure S1B). Principle-component analysis with previous data revealed that mCMG-TBLCs were close to TC1-EGFP cells in PlaB medium after passage 4, while ESC samples from both sets of data clustered together (Figure S1C). To assess whether TBLCs possess the ability to form blastoids, we aggregated 5, 10, 25 and 50 cells of mCMG-TBLCs per microwell of Aggrewell^™^ 400 respectively within the reported medium for generating EPS-blastoid (Li et al., 2019) (Figure 1A and S1D). After 5 days culture, some of the microwells with 5 and 10 cells initiating turned out to be empty because of cell death. Nevertheless, most of the rest cell clumps with different initiating cell numbers started to form a cavity. The cavities grew bigger afterwards to form a blastocyst-like structure in another two days (Figure S1D). We then collected the blastocyst-like structures basic on their morphology and counted their numbers. It turned out that TBLCs gave rise to the highest number of blastocyst-like structures/blastoids with 25 cells initiating. Nearly 80% of microwells formed blastoids (Figure 1B and 1C). The efficiency are much higher than that reported from EPS cells (around 15%) (Li et al., 2019). The average diameter of the blastoids from 25 cells was around 160 μm, slightly bigger than the natural blastocysts (Figure 1D). Previuos work from fusion embryo experiments suggested that the size of blastocysts is dispensable for normal embryo development (Tarkowski, 1961). Hence, we used 25 cells per microwells to initiate the blastoids generation for the rest of our experiments. To investigate the lineage composition of the TBLC-blastoids, we performed immunofluorescence staining for NANOG (marker of epiblast) and CDX2 (marker of TE) or NANOG and GATA6 (marker of PrE) in TBLC-blastoids. It showed that majority of TBLC-blastoids expressed both NANOG, CDX2 and GATA6 (Figure 1E). Yet, we noticed that a small proportion of TBLC-blastoids had no CDX2 positive cells (16%) or have no NANOG positive cells (3%) (Figure S1E). Moreover, while CDX2 positive cells only distributed in the outer layer of blastocysts, 11% TBLC-blastoids contained NANOG negative/CDX2 positive cells in their inner cell clumps (Figure S1F). Excluding these aberrant structures, overall, around 75% TBLC-blastoids expressed all the three lineage markers in a similar pattern with natural blastocysts. To confirm the high efficiency of blastoid generation from TBLCs is independent of specific ES cell line, we derived Tg2a-TBLCs from wild type ESCs, E14Tg2a and generated blastoids with Tg2a-TBLCs (Figure S2A and S2B). It showed that Tg2a-TBLCs form blastoids at a comparable efficiency of mCMG-TBLCs (Figure S2C, S2D and S2E). These results prove that totipotent-like features cells, TBLCs are capable of generating blastoids highly efficiently.

**Figure 1.**
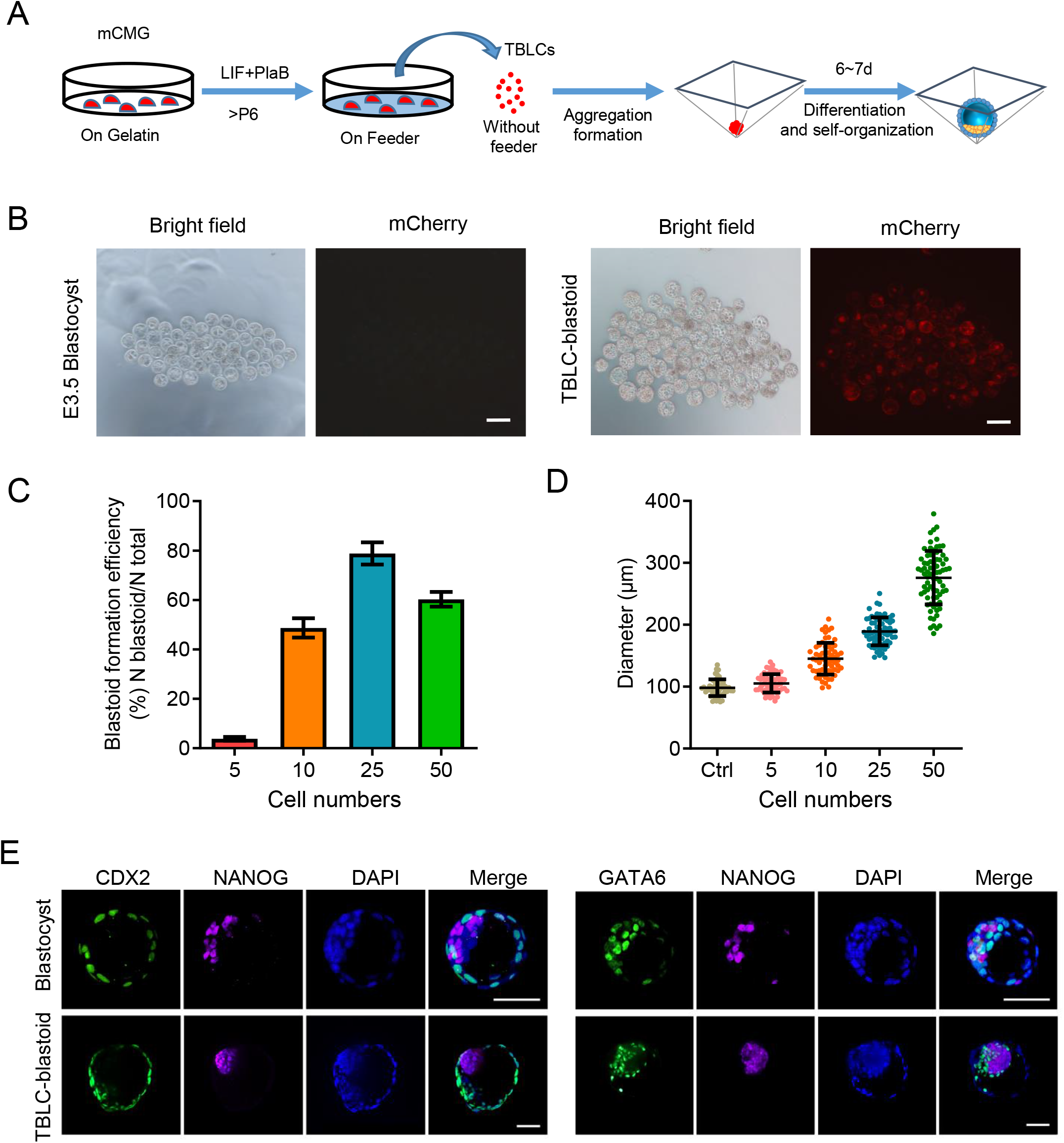
Self-assembly of mouse TBLCs into blastocyst-like Structures (TBLC-Blastoids). (A) Scheme of generating Blastoids from mCMG TBLCs. (B) Microscopy image of blastocysts and TBLC-Blastoids collected from wells with 25-cells per microwell initiation, Scale bar = 200 μm. (C) Quantification of TBLC-blastoids formation efficiency initiating with indicated cell numbers per microwells. Values are means ± SD, n = 3 biological replicates (D) Quantification of the diameter of blastocysts (Ctrl, control) and TBLC-blastoids aggregated from indicated cell numbers per microwells. Values are means ± SD. Each dot represents one blastocyst or blastoid, n = 54, 64, 63, 67, 74 for each group respectively. (E) Immunofluorescence staining of E3.75 blastocysts and TBLC-blastoids for EPI lineage marker (NANOG), TE lineage markers (CDX2) and PE lineage marker (GATA6), Scale bar = 50 μm.

### Transcriptome analysis of blastoids generated from TBLCs

To further assess the transcriptional states of the cells of blastoids from mCMG-TBLCs, we performed 10×Genomics single cell (sc) RNA-seq on total 10068 cells from TBLC-blastoids and compared the data with our scRNA-seq data of natural blastocysts from E3.5 to E4.5 (687 cells in total) and the data of EPS-blastoids (2518 cells in total) (Li et al., 2019). Integrated analysis using SEURAT showed that the cells from TBLC-blastoids, EPS-blastoids and blastocysts largely overlapped with each other (Figure 2A). Followed by using specific lineage marker genes, we identified ICM/epiblast, TE and PrE and an intermediate population in these samples (Figure 2B). Remarkably, cells of TBLC-blastoids expressed all the corresponding lineage marker genes (Figure 2C and Supplementary 3A), though it showed that TE cells from both TBLC and EPS-blastoids expressed lower *Cdx2* and *Eomes*, the two self-renewal regulators of trophoblast cells. Interestingly, consistent with previous report (Posfai et al., 2021), we noted that intermediate population in EPS-blastoids aberrantly expressed mesoderm marker gene, *T* (*Brachyury*) and *Mixl1*. Whereas, those in TBLC-blastoids expressed both ICM and TE marker genes but no T and Mixl1 (Figure 2C). Moreover, TBLC-blastoids showed much less proportion of intermediate cells and more TE cells compared to the EPS-blastoids (Figure 2D). However, the proportion of PrE cells in TBLC-blastoids was less than that of the other two (Figure 2D). Furthermore, unsupervised hierarchical clustering analysis revealed that epiblast and TE lineage of TBLC-blastoids were closer to their counterpart of blastocysts than those of EPS-blastoid (Figure 2E). In summary, these data indicate that TBLC-blastoids have similar lineage composition and cell transcriptome with blastocysts.

**Figure 2.**
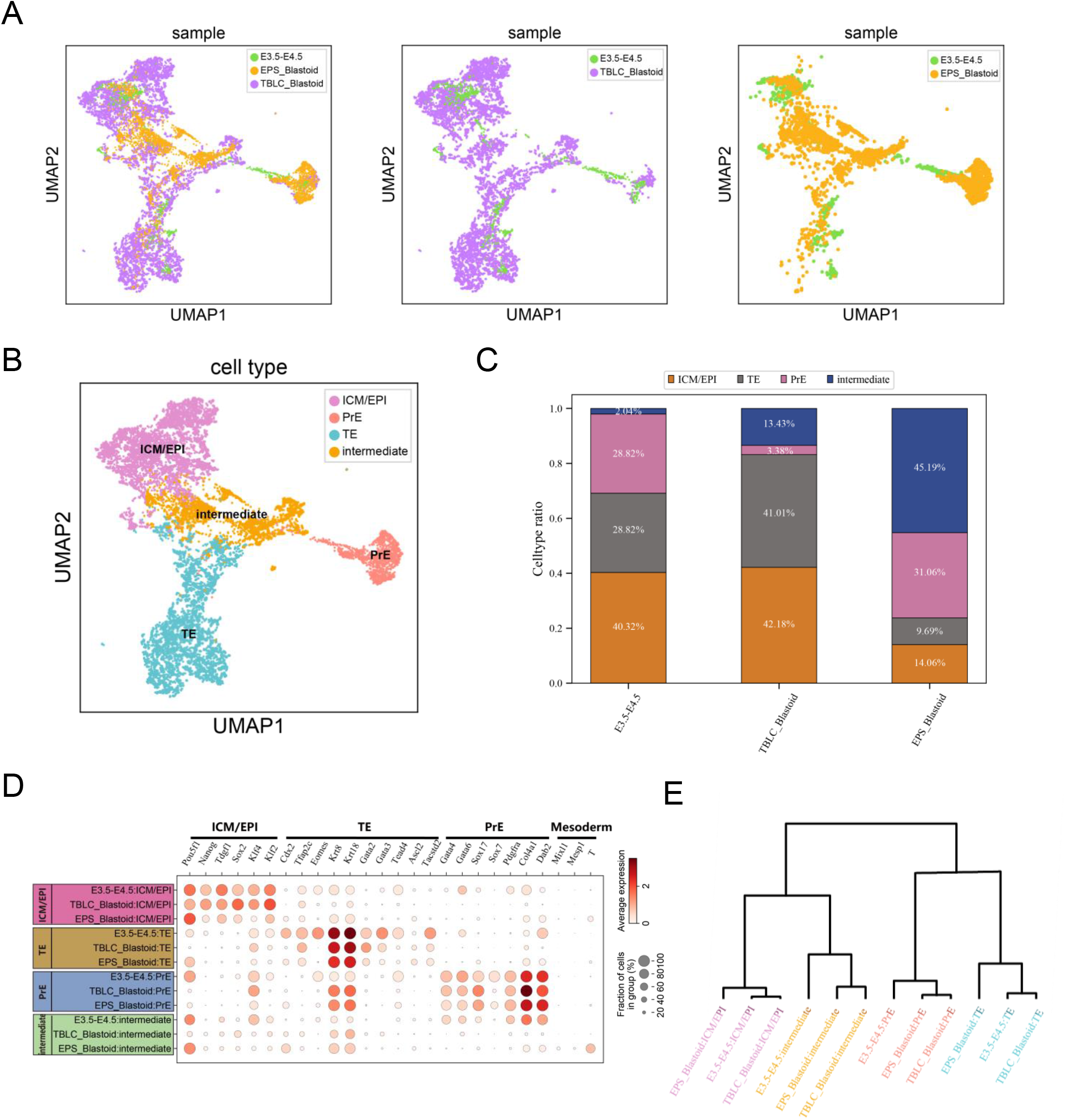
Single-cell transcriptional comparison within TBLC-blastoids, EPS_blastoids and blastocysts. (A) Single-cell UMAP plot of TBLC-blastoids from mCMG (10068 cells), EPS-blastoids (2518 cells) and E3.5-E4.5 embryo cells (687 cells). (B) Single-cell UMAP plot showing the cell types across all samples. The identities of each cluster were determined according to the expression of the lineage markers. (C) The ratio of cell types across all samples. The number within the bar chart showing the percentage of each cell type. (D) Dot plot showing the average expression for selected lineage marker genes of ICM/EPI, PrE, TE and mesoderm. (E) Hierarchical clustering analysis of TBLC-blastoids from mCMG, EPS-blastoids and E3.5-E4.5 embryo cells with cell types in the Figure 2B.

### In vitro and in vivo post-implantation development of TBLC-blastoids

To evaluate whether TBLC-blastoid could grow beyond implantation and recapitulate some aspects of post-implantation development both *in vitro* and *in vivo*, we firstly cultured blastoids from mCMG-TBLCs *in vitro* following the published protocol to transit blastoid/blastocyst to egg-cylinder stage (Figure 3A) (Bedzhov et al., 2014). After cultured in IVC medium for 6 days, comparable propotion of blastoids and blastocysts were attached to the wells. In total, around 10% blastoids and 45% blastocysts formed egg-cylinder structures (Figure S3B). The epiblast (marked by SOX2 or OCT4) of blastoids generated a lumen with the trophectoderm (marked by TFAP2C) on top and surrounded by sporadic GATA6 positive cells (Figure 3B). This indicates that TBLC-blastoids have the ability to develop to egg-cylinder *in vitro*, although with less PrE descendents compared to natural embryos. To further evaluate the postimplantation development potential of TBLC-blastoid *in vivo*, we transplanted day 6 or day 7 TBLC-blastoids into the utero of pseudopregnant mice at 2.5 days post-coitum (dpc) and dissected embryos at E7.5. In total, 16% TBLC-blastoids induced decidualization (Figure 3C, 3D and 3E and S3C). It is noted that most of the deciduae induced by TBLC-blastoids were not fully closure and nearly 95% of them were empty, without embryos inside (Figure 3D). Nevertheless, we found 6 embryo-like structures in 85 TBLC-blastoids induced deciudas. Interestingly, these 6 embryo-like structures were all mCherry-positive. In addition, one of them seems elongated and formed a less-well-organized egg-cylinder structure (Figure 3F). The darker color of the elongated embryo implied that this embryo was absorbing. Yet, immunofluorescence analysis revealed that there were a few SOX2 and TFAP2C positive cells in TBLC-blastoids derived embryo (Figure 3G). These results indicate that TBLC-blastoid can develop beyond the implantation *in vitro* and induce deciduae *in vivo*, however, they are unable to develop normally *in vivo*.

**Figure 3.**
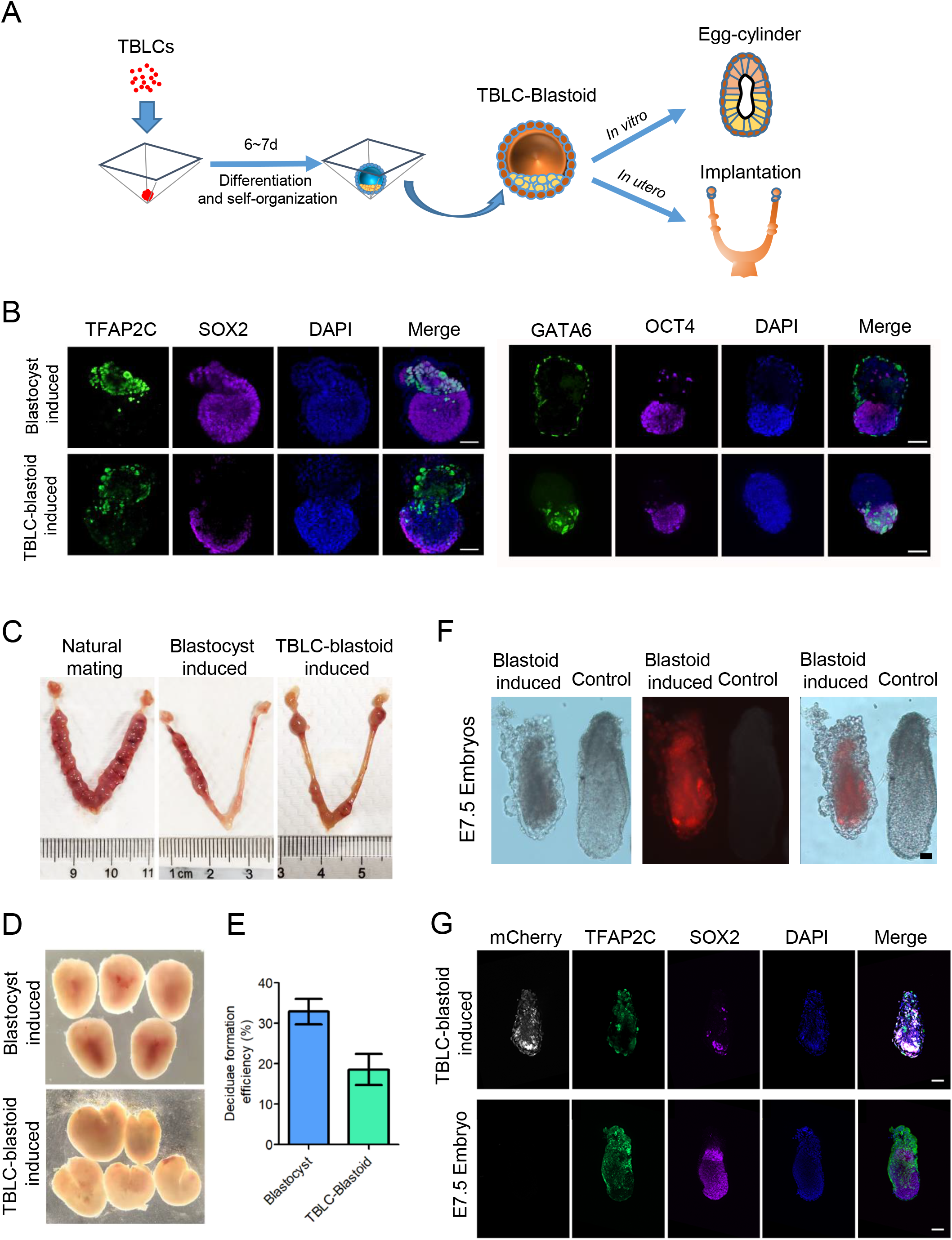
Post-implantation development *in vitro* and *in vivo* of TBLC-blastoids. (A) Schematic of *in vitro* and *in vivo* post-implantation development assessment of TBLC-blastoids. (B) Immunofluorescence staining of blastocyst (top) and TBLCblastoid (bottom)-derived post-implantation egg-cylinder-like structures for TFAP2C and SOX2 (left) or GATA6 and OCT4 (right), Scale bar = 200 μm. (C) Bright-field image of mouse uterus with deciduae at E7.5 induced from natural mating or transferred blastocysts/blastoids (D) Brightfield images of deciduae dissectedfrom 7.5 dpc mice (Control) or surrogate mice at 7.5 dpc with TBLC-blastoids transferred at 2.5 dpc. (E) Quantification of the efficiency of decidua formation from blastocysts or blastoids after transferred into mice uterus (date in Figure S3C) (F) Brightfield and fluorescent images of E7.5 embryos (Control) and TBLC-blastoid derived structures recovered from deciduae, Scale bar = 100 μm. (G) Whole-mount immunofluorescence staining for mCherry, TFAP2C, and SOX2 of E7.5 embryo and a embryo-like structure recovered from TBLC-blastoids induced decidua at 7.5 dpc. Scale bar = 500 μm.

## Discussion

Generation of blastocyst-like structures *in vitro* from totipotent-like feature cells supplies a readily accessible, scalable, tractable and highly malleable system to study pre-implantation and early post-implantation embryogenesis. However, the efficiency of previous reported mouse blastoids generation from EPS is low and a considerable population of mesoderm-like cells exist in EPS-blastoids. In this work, we provide the evidence that TBLCs, a recently reported totipotent-like cell type, are capable of forming blastoids highly efficiently (around 80%) when aggregated in a 3D differentiation system *in vitro*.

Both immunostaining of lineage specific protein and scRNA-seq confirmed that TBLC-blastoid contain the three early embryonic lineages, ICM/Epi, TE and PrE. TBLC-blastoids contain fewer intermediate cells compared to blastoids from EPS cells (around 10% *versus* 45%) and intermediate cells show no expression of mesodermal markers. Furthermore, TBLC-blastoids contained much higher portion of TE cells than EPS-blastoids and both epiblast and TE lineage of TBLC-blastoids are closer with their counterpart of blastocysts than that of EPS-blastoids. In addition, high efficiency of generation blastoids from TBLCs could supply an improved system for high-throughput screening the effects of gene mutations, medicine or toxins for early embryogenesis. Moreover, it can be used to screen cytokines or small inhibitors to generate fully functional synthetic blastocyst.

Hitherto, there were no full-term embryos derived from blastoids reported. Both our and others work showed that epiblast cells of blastoids recapitulating early post-implantation development process *in vitro* but not *in vivo* suggested that the implantation process *in vivo* was not well established although deciduae were induced. Indeed, we noticed that the deciduae from blastoids were fragile and not fully closure. This is probably because of the abnormal extraembryonic cell lineages of blastoids. Our data revealed that TBLC-blastoid contains less PrE population compared to nature blastocyst (3% *versus* 28%). Moreover, while ICM/epiblast and TE cells of TBLC-blastoid are close to that of blastocyst, PrE cells of TBLC-blastoid are close to that of EPS-blastoid. In addition, low expression of *Cdx2* and *Eomes* in TE cells of blastoids implies that polar TE cells in blastoids might be insufficient (Frias-Aldeguer et al., 2020). Further studies to modify the medium to obtain proper proportion faithful PrE and TE cells in blastoids are needed.

## METHODS AND MATERIALS

### Cell culture

mES culture: mES cell lines were cultured in mES medium [DMEM supplemented with 15% Fetal Bovine Serum (FBS, 16000-044), 1x NEAA (11140-050), 1x GlutaMAX (35050-061), 1x Sodium pyruvate (11360-070), 1 U/mL penicillin and 1 mg/mL streptomycin (Gibco, 15070063), 0.1 mM β-mercaptoethanol (Sigma, 63689), and 10^3^ U/mL LIF (Merck, ESG1107)] on gelatin under 5% CO_2_ at 37□.

TBLC culture: To generate TBLCs (totipotent blastomere-like cells), the mES cells were seeded on a layer of irradiated ICR mouse embryonic fibroblasts (MEF) and culture in mES medium (see above) supplemented with 2.5 nM PlaB (Tocris, 6070). Cells were passaged every 3-4 days with 0.05% trypsin-EDTA and seeded on MEF feeder layers at split ratios between 1:3 to 1:5. After 6 passages, TBLCs were used for further experiments.

### Fluorescence activated Cell Sorting (FACS)

Cells were digested into single cells using 0.05% trypsin-EDTA and resuspended in FACS medium (DPBS with 5% KSR). Cell suspension was then filtered through 40 μm cell strainers (Falcon, 352340). Next, the mCherry-positive TBLCs were sorted on a BD FACSAria Fusion. Data analysis was performed using FlowJo 10.

### Blastocyst-like Structure Generation on AggreWells

AggreWell 400 (STEMCELL Technologies, 34415) was prepared following the manufacturer’s instructions. EPS blastoid basal medium is composed of 25% TSC basal medium [RPMI 1640 (11875-093) supplemented with 20% FBS (16000-044), 1% GlutaMAX (35050-061), 1% Sodium pyruvate (11360-070), and 0.1 mM 2-mercaptoethanol (21985-023), all from Thermo Fisher Scientific], 25% N2B27 basal medium [1:1 mixture of DMEM/F-12 (11330-032) and Neurobasal (21103-049) supplemented with 0.5% N2 (17502-048), 0.5% B27 (17504-044), 1% NEAA (11140-050), 1% GlutaMAX (35050-061), 0.1 mM 2-mercaptoethanol (21985-023), and 5% KnockOut Serum Replacement (Optional, 10828-028), all from Thermo Fisher Scientific)], and 50% M16 (Sigma, M7292).The corresponding number sorted TBLCs were resuspended in EPS-blastoid basal medium supplemented with 2 μM ROCK inhibitor Y-27632 (Reagents Direct, 53-B80-50), 12.5 ng/mL rhFGF4 (R&D, 235F4025), 0.5 μg/mL Heparin (Sigma-Aldrich, H3149), 3 μM CHIR99021 (Reagents Direct, 27-H76), 5 ng/mL BMP4 (Proteintech, HZ-1040), and 0.5 μM A83-01 (Axon Medchem, 1421) and seeded into one well of the 24-well AggreWell plate. After mixed, the plate was centrifuged for 1 min at 200 g and then placed at 37□ and 5% CO_2_. The day of cell seeding was counted as day 0 of the process. Medium was half removed 24 h later (day 1) and replaced with fresh medium without Y-27632 every other day.

### In Vitro Culture of Blastocysts and Blastocysts-like Structures

Blastocysts and blastocysts-like structures were cultured in vitro according to the protocol published previously (Bedzhov et al., 2014). Blastocysts were cultured in KSOM medium (Millipore, MR-107-D) until E4.5, and then were transferred into the acidic Tyrode’s solution as the zona pellucida gradually disappear. The zona-free blastocysts and the generated blastocysts-like structures were washed twice with preequilibrated IVC1 medium, and transferred in to a μ-Silde 8-well (ibidi, 80826), containing pre-equilibrated IVC1 medium, about 4-8 blastocysts or 10 blastoids per well, cultured in incubators at 37°C, 5% O_2_ and 5% CO_2_. After 3 days, part of blastocysts or blastoids attached to the plate, the medium was switched to IVC2 medium. After 2-4 days, the structures were fixed with 4% PFA for 20 min at room temperature for immunofluorescence staining. IVC1 medium consisted of advanced DMEM/F12 medium (Gibco, 11320033), 20% FBS, 2 mM L-glutamine, 1% penicilin-streptomycin, 1% 1%ITS-X (Gibco, 51500056), 8 nM β-estradiol (Sigma, E2758), 200 ng/mL progesterone (Selleck, S1705), 25 μM N-acetyl-L-cysteine (Selleck, S1623). IVC2 medium consisted of advanced DMEM/F12 medium, 30% KSR, 2 mM L-glutamine, 1% penicilin-streptomycin, 1% ITS-X, 8 nM β-estradiol, 200 ng/mL progesterone, 25 μM N-acetyl-L-cysteine.

### Immunostaining

E3.5 blastocysts and blastocyst-like structures, were fixed in 4% paraformaldehyde (PFA, Meilunbio, MA0192) for 20 min at room temperature. Natural embryos or blastoids were fixed in 4% PFA overnight at 4°C.The fixed samples were washed three times for 10 min in PBST [DPBS (HyClone, SH30028.02) with 0.1% Triton-X100 (Macklin, T824275)], permeabilized for 30 min at room temperature in DPBS/0.3% Triton-X100 and block by blocking buffer (PBST containing 3% donkey serum (Sigma, D9663) and 1% BSA (Sigma, V900933) for 2 h. Primary antibody incubation was performed overnight at 4 □ in blocking buffer. The following day, embryos were washed three times for 10 min in PBST, then incubated with secondary antibody (1:500) in blocking buffer 4 h at room temperature at 4□. After finished the incubation, embryos were washed and stained with DAPI (Sigma, D9542) for 5 min. Imaging occurred on a Carl Zeiss LSM800 confocal microscope (Carl Zeiss, LSM800) and images were processed using ZEN software (Carl Zeiss). The antibodies used during the experiment are listed in Supplemental Information (Supplementary Table 1).

### Blastoid Transfer

All procedures related to animals were performed following the ethical guidelines of the Guangzhou Laboratory. Blastoids were manually picked up under a stereomicroscope and transferred into M2 droplets using a mouth pipette. The surrogate at 2.5 days post coitum (dpc) was anesthetized with 2,2,2-Tribromoethanol (Sigma, T48402) and tert-Amyl alcohol (Sigma, 152463) and the uterine horn was exposed surgically. After three washes in M2, Blastoids were transferred to the uterine horn, which was previously punctured with a 27G needle. Around 20 Blastoids were transferred into each uterine horn. At 7.5 dpc, deciduae were dissected out of the uterus, and embryo-like structures were dissected out of the deciduae. Tissue samples were fixed with 4% PFA for immunostaining.

### Bioinformatics analysis

Bulk RNA-seq analysis: The quality of raw sequencing reads were performed using Fastqc (v0.11.2). Sequencing adapters and low-quality reads/bases were further removed using trim_galore (v0.6.4). Clean reads were then aligned to the mouse reference genome (GRCm38) using STAR (v2.7.6a) (Dobin et al., 2013) and only uniquely mapped reads were retained. Raw counts for each gene were estimated using RSEM (v1.3.1) from the aligned bam files. The published RNA-seq data set from Shen et al., (Shen et al., 2021) was downloaded from the NCBI Gene Expression Omnibus (GEO accession number GSE168728). Batch correction for our RNA-seq data set and Shen et al. data set was perfromed by using the Combat-seq (Zhang et al., 2020). Principle components analysis was performed using the R function Prcomp. Heatmap was plotted using pheatmap package. To identify pluripotent and totipotent genes compared with mESCs, differentially expressed gene analysis was performed using R package DESeq2 (adjusted p value < 0.05 and fold change > 2) (Love et al., 2014).

Single-cell RNA-seq data processing: Single-cell RNA-seq reads were first mapped to the mouse reference genome (mm10) using STAR(v2.7.6a). Next, we filtered the cells with a minimum cut-off of 3000 expressed genes per cell. After filtering, a total of 13273 cells were retained for the down-stream analyses. Then, we integrated single-cell data using Seurat’s canonical correlation analysis (CCA) integration tool (Seurat v4.0.4) with FindIntegrationAnchors and IntegrateData; Scale data with 3000 highly variable genes were used to compute PCA, 30 principal components were used to calculate UMAP coordinates with RunUMAP. Cell clusters were identified by a shared nearest neighbor (SNN) modularity optimization based on clustering algorithm with FindClusters function. The marker genes expression on UMAP embedding and dot plot were plotted using the log-normalized data. EPS_blastoids gene expression matrices were downloaded as provided by Li et al (GSE135701).

### Materials availability

All unique/stable reagents generated in this study are available from the Lead Contact with a completed Materials Transfer Agreement.

### Data availability1

All raw sequencing data can be accessed at NCBI Gene Expression Omnibus (GEO). Raw count of scRNA-sequencing data of E3.5-E4.5 blastocyst (in-house data) can be found in supplement table 2.

## ACKNOWLEDGMENTS

We thank Dr. Peng Du (Peking University) for their TC1-EGFP ESCs and TC1-EGFP TBLCs, thank Dr. Jiekai Chen and Duanqing Pei for their single cell RNA seq data of blastocysts.

## AUTHOR CONTRIBUTIONS

M.Z. conceived, designed and conducted the studies; P.Z., B.H., S.S., W.W., performed experiments; X.Z., performed bioinformatics analysis; M.Z., P.Z. and X.Z. wrote the paper, with input from all authors.

---

**Figure S1.**
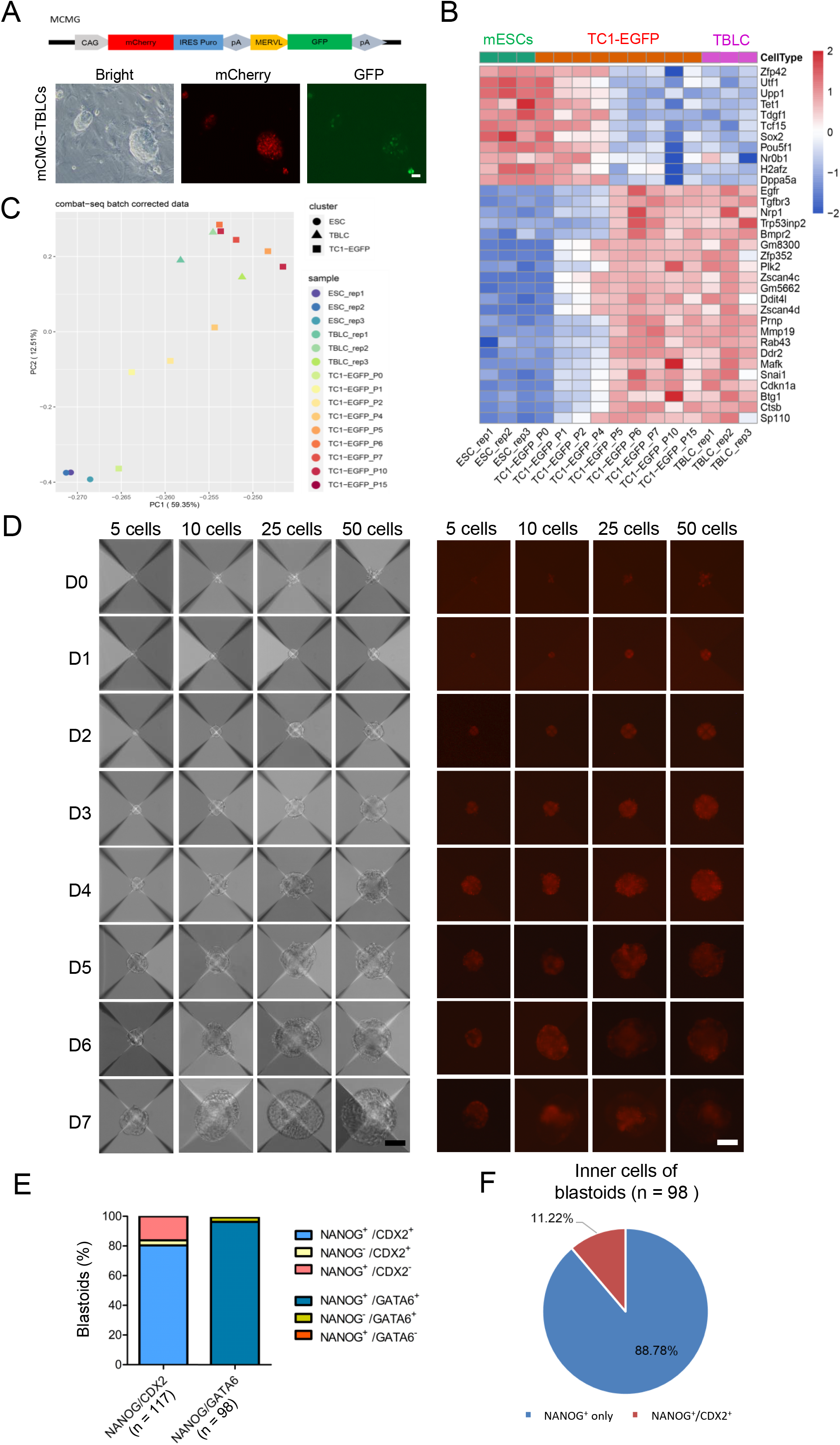
Self-assembly of mCMG-TBLCs (induced from mCMG cell line) into blastocyst-like structures, Related to Figure 1. (A) Diagram of CAG::mCherry; MERVL::GFP (mCMG) transgenic ES cell line (upper) and representative phase-contrast and fluorescence images of TBLC induced from mCMG cell line, Scale bar = 50 μm. (B) Heatmap of the relative expression of representative pluripotent and totipotent genes in mESCs and TBLCs, co-analysis with the data from GSE168728 (Shen et al., 2021) (C) Transcriptome-based PCA of our mCMG-TBLCs (TBLC_ rep1/2/3) and different passages TC1-EGFP cells cultured in the ES medium with PlanB reported by Hui et al. 2021. (D) Representative phase-contrast (left) and fluorescence (right) images of aggregates in the indicated days showing the formation of mCMG TBLC-Blastoids initiating with 5-, 10-, 25- or 50-cells/microwell, Scale bar = 50 μm. (E) Distribution percentage of TBLC-blastoids expressed indicated types of NANOG/CDX2 and Nanog/ GATA6. (F) Proportion of TBLC-blastoids with NANOG positive cells only or both NANOG positive and CDX2 positive cells in their inner cell clumps.

**Figure S2.**
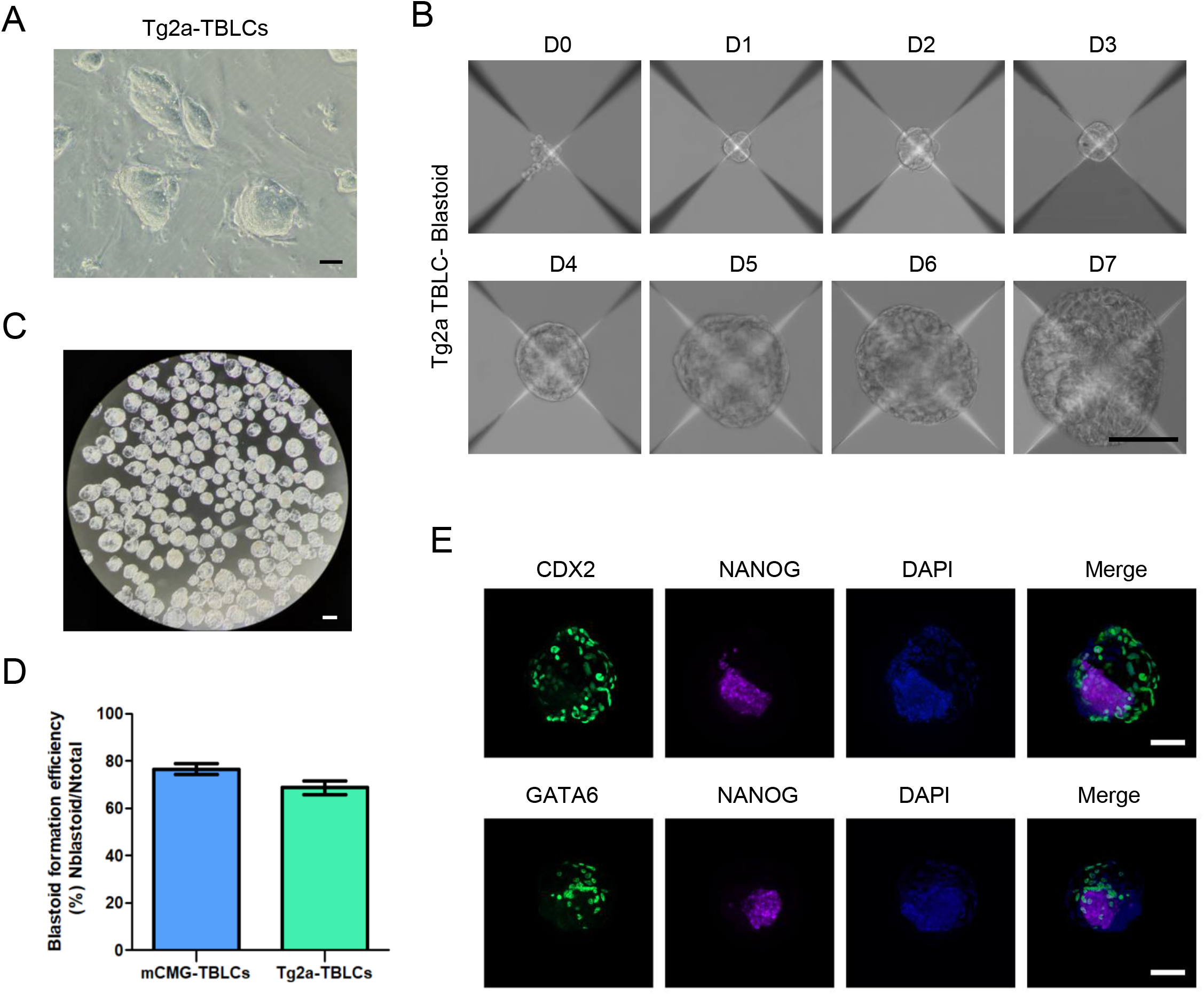
Self-assembly of Tg2a-TBLCs (induced from E14Tg2a cell line) into Blastocyst-like Structures. Related to Figure 1. (A) Representative phase-contrast images of Tg2a-TBLCs induced from E14Tg2a cell line, Scale bar = 50 μm. (B) Representative phase-contrast images of aggregates at the indicated days showing the formation of Tg2a TBLC-blastoids initiating with 25 cells per microwell, Scale bar = 50 μm. (C) Brightfield image of Tg2a-TBLC-blastoids collected from microwells at day 7 of aggregation, Scale bar = 200 μm. (D) Quantification of Tg2a TBLC-Blastoids formation efficiency (n=10 independent assays for each TBLC cell line) initiating with 25 cells per microwell. (E) Immunofluorescence staining of Tg2a-TBLC-blastoids for EPI lineage marker (NANOG), TE lineage markers (CDX2) and PE lineage marker (GATA6), Scale bar = 50 μm.

**Figure S3.**
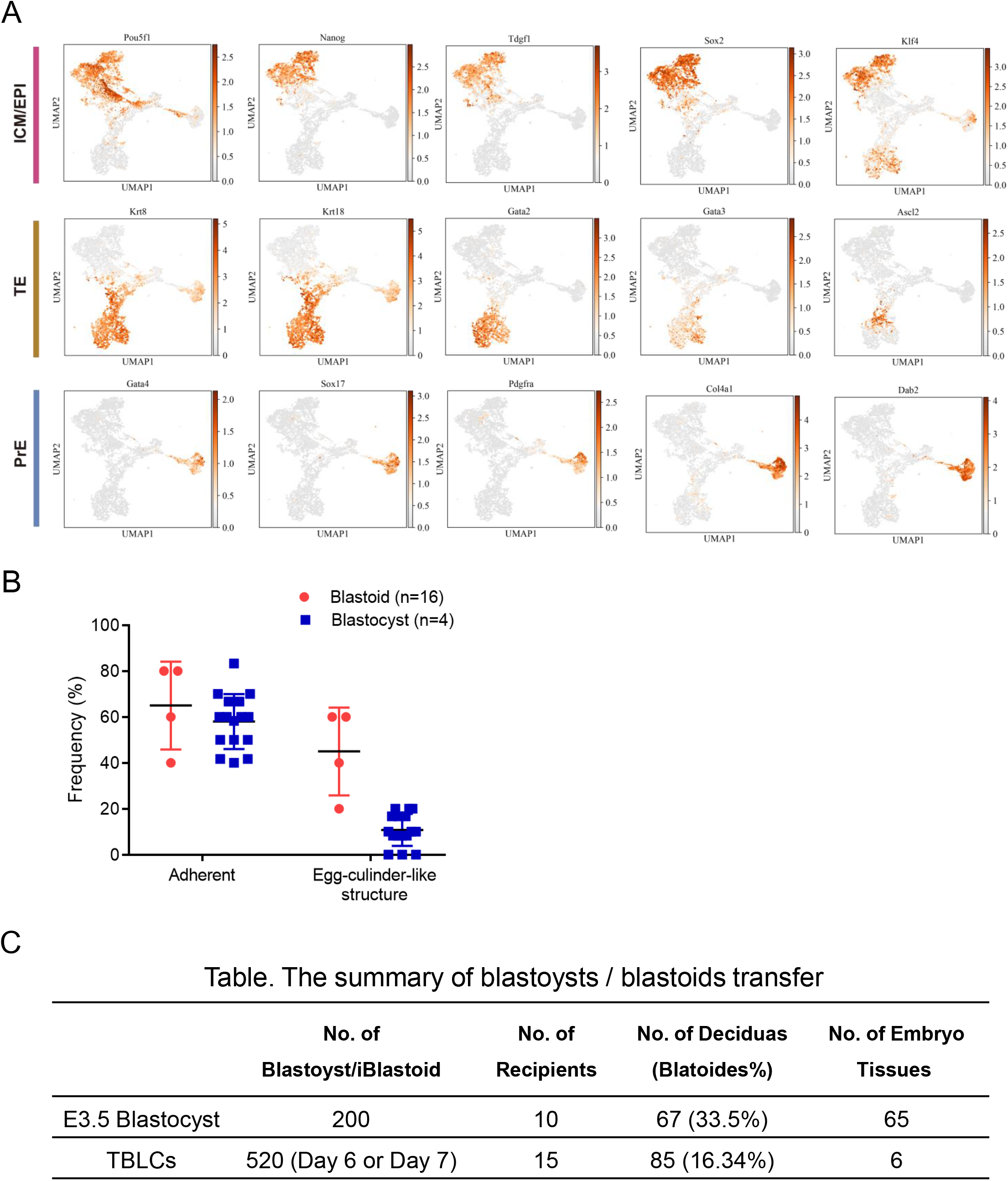
Related to Figure 2 and Figure 3. (A) Visualization of marker genes of ICM/EPI, PE and TE in the UMAP. (B) Quantification of the percentage of adherent embryos and formed postimplantation embryo-like structures after in vitro culture of blastocysts and TBLC-blastoids in each well, n =4 for blastocysts and 16 for blastoids, data from 2 independent experiments. (C) Table to summary the blastoysts / blastoids transfer.

