## Supplemental Data 1 for "Highly efficient generation of blastocyst-like structures from mouse totipotent blastomere-like cells"

Supplementary Table 1. Antibodies Used In This Study

| Antibodies | SOURCE | IDENTIFIER | Dilution |
| --- | --- | --- | --- |
| rabbit-anti-NANOG | Bethyl | Cat# A300-397A-M | 1:200 |
| goat-anti-GATA6 | R&D | Cat# AF1700 | 1:200 |
| Mouse anti-AP2γ (TFAP2C) | Santa Cruz | Cat# sc-12762 | 1:100 |
| Rabbit anti-SOX2 | CST | Cat# 23064S | 1:200 |
| mouse-anti-OCT4 | Santa Cruz | Cat# sc-5279 | 1:100 |
| mouse-anti-CDX2 | Abcam | Cat# ab76541 | 1:200 |
| Anti-mCherry antibody | Abcam | Cat# ab167453 | 1:200 |
| Donkey anti-Goat IgG (H+L) Highly Cross-Adsorbed, Alexa Fluor Plus 488 | Invitrogen | Cat# A32814 | 1:500 |
| Donkey anti-Rabbit IgG (H+L) ReadyProbes™, Alexa Fluor 488 | Invitrogen | Cat# R37118 | 1:500 |
| Donkey anti-Mouse IgG (H+L) Highly Cross-Adsorbed, Alexa Fluor 647 | Invitrogen | Cat# A31571 | 1:500 |
| Donkey anti-Rabbit IgG (H+L) Highly Cross-Adsorbed, Alexa Fluor 647 | Invitrogen | Cat# A31573 | 1:500 |
| Donkey anti-Mouse IgG (H+L) Highly Cross-Adsorbed, Alexa Fluor 488 | Invitrogen | Cat# A32766 | 1:500 |
